# Genome-wide Prediction of DNase I Hypersensitivity Using Gene Expression

**DOI:** 10.1101/035808

**Authors:** Weiqiang Zhou, Ben Sherwood, Zhicheng Ji, Fang Du, Jiawei Bai, Hongkai Ji

## Abstract

We evaluate the feasibility of using a biological sample’s transcriptome to predict its genome-wide regulatory element activities measured by DNase I hypersensitivity (DH). We develop BIRD, Big Data Regression for predicting DH, to handle this high-dimensional problem. Applying BIRD to the Encyclopedia of DNA Element (ENCODE) data, we found that gene expression to a large extent predicts DH, and information useful for prediction is contained in the whole transcriptome rather than limited to a regulatory element’s neighboring genes. We show that the predicted DH predicts transcription factor binding sites (TFBSs), prediction models trained using ENCODE data can be applied to gene expression samples in Gene Expression Omnibus (GEO) to predict regulome, and one can use predictions as pseudo-replicates to improve the analysis of high-throughput regulome profiling data. Besides improving our understanding of the regulome-transcriptome relationship, this study suggests that transcriptome-based prediction can provide a useful new approach for regulome mapping.

## INTRODUCTION

A fundamental question in functional genomics is how genes’ activities are controlled temporally and spatially. To answer this question, it is crucial to comprehensively map activities of all genomic regulatory elements (i.e., regulome) and understand the complex interplay between the regulome and transcriptome (i.e., transcriptional activities of all genes). Regulome mapping has been accelerated by high-throughput technologies such as chromatin immunoprecipitation coupled with high-throughput sequencing (Johnson et al. 2007) (ChIP-seq) and sequencing of chromatin accessibility (e.g., DNase-seq (Crawford et al. 2006) for DNase I hypersensitivity, FAIRE-seq (Giresi et al. 2007) for Formaldehyde-Assisted Isolation of Regulatory Elements, and ATAC-seq (Buenrostro et al. 2013) for Assaying Transposase-Accessible Chromatin). So far these technologies have only been applied to interrogate a small subset of all possible biological contexts defined by different combinations of cell or tissue type, disease state, time, environmental stimuli, and other factors. A major limitation of current high-throughput technologies is the difficulty to simultaneously analyze a large number of different biological contexts. This limitation along with various practical constraints such as lack of materials, antibodies, resources, or expertise has hindered their application by the vast majority of biomedical investigators from small laboratories.

For the study of regulome-transcriptome relationship, numerous researchers have examined how genes’ transcriptional activities can be predicted using activities of their associated regulatory elements (Natarajan et al. 2012; Cheng et al. 2012; Kumar et al. 2013). However, the interplay between regulome and transcriptome is bidirectional due to presence of feedback (Neph et al. 2012; Voss and Hager 2014). A systematic understanding of this relationship in the reverse direction – to what extent regulatory elements’ activities can be predicted by transcriptome – is still lacking. We investigate this reverse prediction problem using DNase I hypersensitivity (DH) and gene expression data generated by the Encyclopedia of DNA Elements (ENCODE) Project (ENCODE Project Consortium 2012). Besides creating a more complete picture of the regulome-transcriptome relationship, this investigation also has important practical implications for regulome mapping. Gene expression is the most widely measured data type in high-throughput functional genomics. Measuring expression does not require large amounts of materials and complex protocols, and technologies for expression profiling are relatively mature. As a result, expression data are routinely collected even when other functional genomic data types are difficult to generate due to technical or resource constraints. Today, the Gene Expression Omnibus (GEO) database (Edgar et al. 2002) contains 200,000+ human gene expression samples from a broad spectrum of biological contexts, as compared to only ~7000 human ChIP-seq, DNase-seq, FAIRE-seq and ATAC-seq samples available in GEO. We reasoned that if one can use the ENCODE data to build prediction models and apply these models to existing and new transcriptome data to predict regulome, the catalog of biological contexts with regulome information may be quickly expanded (**Fig. 1a**). This will provide a useful approach for regulome mapping that is complementary to existing experimental methods. It will also allow researchers to more effectively use expression data to study gene regulation. Unlike a recent study that imputes one functional genomic data type based on multiple other data types which are non-trivial to collect (Ernst and Kellis 2015), prediction in this study is based on one single but widely available data type and hence can have a substantially broader range of applications. During our investigation, we develop a big data regression approach, BIRD, to handle the prediction problem where both predictors (i.e., transcriptome) and responses (i.e., regulome) are ultra-high-dimensional, which is an emerging problem in the analysis of big data.

**Figure 1.**
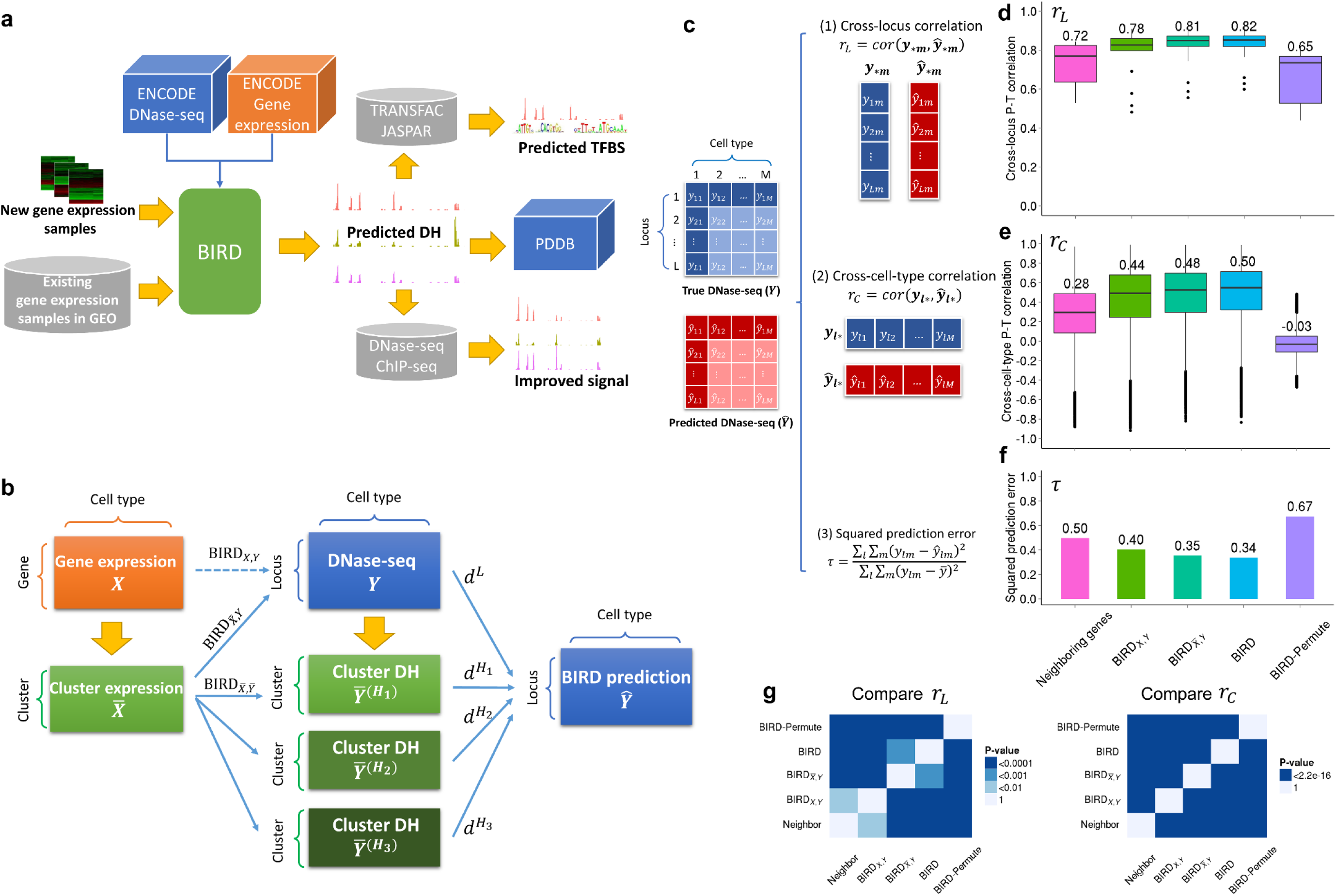
BIRD – concepts and evaluation. (**a**) Outline of the study. ENCODE DNase-seq and exon array data are used to train BIRD. Users can apply BIRD to new or existing gene expression samples to predict DH. The predicted DH can be used to predict TFBSs, convert expression samples in GEO into a regulome database (PDDB), and improve DNase-seq and ChIP-seq data analyses. (**b**) Overview of BIRD. Instead of using expression of individual genes as predictors to predict DH at each locus (BIRD_X,Y_), BIRD first groups co-expressed genes into clusters (i.e., gene-cluster) and uses the clusters’ mean expression levels as predictors to predict the DH level at each genomic locus 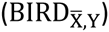. Additionally, BIRD also groups correlated loci (i.e., loci with co-varying DH) into different levels of clusters (i.e., DHS-cluster) and predicts the DH level for each cluster 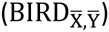. Finally, BIRD combines the locus-level and cluster-level predictions via model averaging. (**c**) Statistics used to evaluate prediction accuracy. (**d**) Cross-locus P-T correlation (*r_L_*) for different methods. For each method, the boxplot and number show the distribution and mean of *r_L_* from the 17 test cell types. (**e**) Cross-cell-type P-T correlation (*r_C_*) for different methods. For each method, the distribution and mean of *r_C_* from the 912,886 genomic loci are shown. (**f**) Squared prediction error (*τ*) for different methods. (**g**) Two-sided Wilcoxon signed-rank test p-values for comparing prediction performance (*r_L_* or *r_C_*) of different methods. We did not perform similar test for the squared prediction error (*τ*) since there is only one *τ* for each method.

## RESULTS

### Big Data Regression for Predicting DNase I hypersensitivity (BIRD)

We obtained DNase-seq and exon array (i.e., gene expression) data for 57 distinct human cell types with normal karyotype from ENCODE (**Supplementary Table 1**). The 57 cell types were randomly partitioned into a training dataset (40 cell types) and a test dataset (17 cell types). After filtering out genomic regions with weak or no DH signal across all 40 training cell types, 912,886 genomic loci (also referred to as “DNase I hypersensitive sites” or “DHSs”) with unambiguous DNase-seq signal in at least one training cell type were retained for subsequent analyses (**Methods**).

Our goal is to use gene expression to predict DH. This can be formulated as a problem of fitting millions of regression models, one per genomic locus, to describe the relationship between DH (response) and gene expression (predictor). The regression for each locus can be constructed using either its neighboring genes or all genes as predictors (**Supplementary Fig. 1**). We tested both strategies (see **Methods and Supplementary Figs. 2-4** for details). The latter strategy requires dealing with a challenging big data regression problem which involves fitting about 1 million high-dimensional regressions, each with a large number (18,000+) of predictors and small sample size. To cope with the high dimensionality and heavy computation, we developed the BIRD algorithm (**Fig. 1b, Methods**). The elementary BIRD model 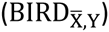 groups correlated predictors into clusters and transforms each cluster into one predictor. A prediction model for each genomic locus is then constructed using the transformed predictors. Clustering reduces the predictor dimension, mitigates co-linearity, makes the predictors less sensitive to measurement noise, and improves prediction accuracy. A variant of the elementary model, 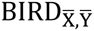, further clusters co-activated DHSs (i.e., correlated responses) and predicts the mean DH level of each cluster. Finally, the compound BIRD model (BIRD) integrates the locus-level predictions from 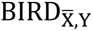 and the cluster-level predictions from 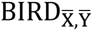 via model averaging to better balance the prediction bias and variance. A systematic benchmark analysis shows that BIRD not only offers the computational efficiency suitable for big data regression, but also had the best prediction performance in our problem compared to other methods (**Supplementary Methods, Supplementary Fig. 4**).

### Predicting DNase I Hypersensitivity Based on Gene Expression

We applied BIRD to the 40 training cell types to build prediction models, and evaluated their prediction performance in the 17 test cell types using three types of statistics: (1) the Pearson correlation between the predicted and true DH values (or P-T correlation) across different genomic loci within each cell type (*r_L_*), (2) the P-T correlation across different cell types at each genomic locus (*r_C_*), and (3) the total squared prediction error scaled by the total DH data variance (*τ*) (**Fig. 1c, Methods**). These analyses led to the following findings.

*Gene expression provides valuable information for predicting DH*. **Figure 1d-g** compares *r_L_*, *r_C_* and *τ* from different methods (**Methods**). These plots show that the elementary BIRD model 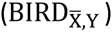 significantly increased the P-T correlation (*r_L_* and *r_C_*) and substantially decreased the squared prediction error (*τ*) compared to random prediction models (BIRD-Permute). The random prediction models were obtained by applying the same 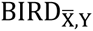 analysis after permuting the link between DNase-seq and gene expression data in the training dataset.

*Prediction based on the whole transcriptome substantially improves prediction based on a genomic locus’ neighboring genes*. We tested the neighboring gene approach by gradually increasing the number of neighboring genes and identified the optimal performance (**Methods, Supplementary Fig. 2**). Compared to the best prediction performance by the neighboring gene approach, 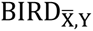 substantially increased the prediction accuracy (**Fig. 1d-g**), indicating that not all information useful for prediction is contained in neighboring genes. This is plausible biologically because DH of a locus may be correlated *in trans* with expression of TFs that bind to the locus, genes that co-express with these TFs, and genes that co-express with the target gene controlled *in cis* by the locus. Moreover, since cell-type-specific transcription of a gene may be controlled by multiple *cis*-regulatory elements, DH of a particular regulatory element may not always correlate well with its neighboring gene expression.

*Clustering correlated predictors (i.e., co-expressed genes) helps prediction*. In 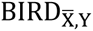, correlated predictors are consolidated by clustering. BIRD_X,Y_ is a special case of 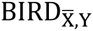 in which predictors are not clustered whereas all subsequent predictor selection and model fitting procedures remain the same (**Methods**). Compared to BIRD_X,Y_, 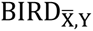 produced higher prediction accuracy (**Fig. 1d-g**, **Supplementary Fig. 3b**). This shows that in a high-dimensional regression setting where predictors far outnumber the sample size, clustering correlated predictors before variable selection and model fitting, a technique not widely used in high-dimensional regression literature, can improve the model compared to conventional techniques (Tibshirani 1996; Fan and Lv 2008) that directly apply variable selection to reduce the predictor dimension.

*DH variation across different genomic loci within a cell type can be accurately predicted*. In the 17 test cell types, the mean cross-locus P-T correlation *r_L_* of 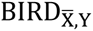 was 0.81 (**Fig. 1d**). Interestingly, random prediction models were also able to produce large *r_L_* (**Fig. 1d**, mean = 0.65). This is because different loci have different DH propensity, consistent with observations in a previous study (Ernst and Kellis 2015). For instance, some loci tend to show higher DH signal than other loci in most cell types (**Supplementary Fig. 5**). As a result, using the average DH profile of all training cell types can predict the cross-locus DH variation in a new cell type with good accuracy (Ernst and Kellis 2015), even though such predictions are cell-type-independent and remain the same for all new cell types. Our random prediction models were generated by permutations that did not perturb the locus-specific DH propensity. Therefore, their *r_L_* was large. Since 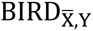 uses cell-type-dependent information carried by transcriptome, its predictions are more accurate (**Fig. 1d**).

*DH variation across cell type can be predicted, although it is more challenging than predicting cross-locus variation*. **Figure 2a** shows an example demonstrating that the true cross-cell-type DH variation measured by DNase-seq can be captured by BIRD predictions, but not by the mean DH profile of all training cell types. Comparing the cross-locus P-T correlation (*r_L_*) in **Figure 1d** with the cross-cell-type P-T correlation (*r_C_*) in **Figure 1e**, *r_L_* on average was substantially larger than *r_C_* (e.g., 0.81 vs. 0.48 for 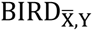. Unlike *r_L_*, the distribution of *r_C_* for random prediction models was centered around zero (**Fig. 1e**, mean = −0.03) because the cross-cell-type prediction accuracy was evaluated within each locus and hence not affected by locus effects. Compared to random prediction models, 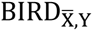 significantly increased *r_C_* (**Fig. 1e,g**).

**Figure 2.**
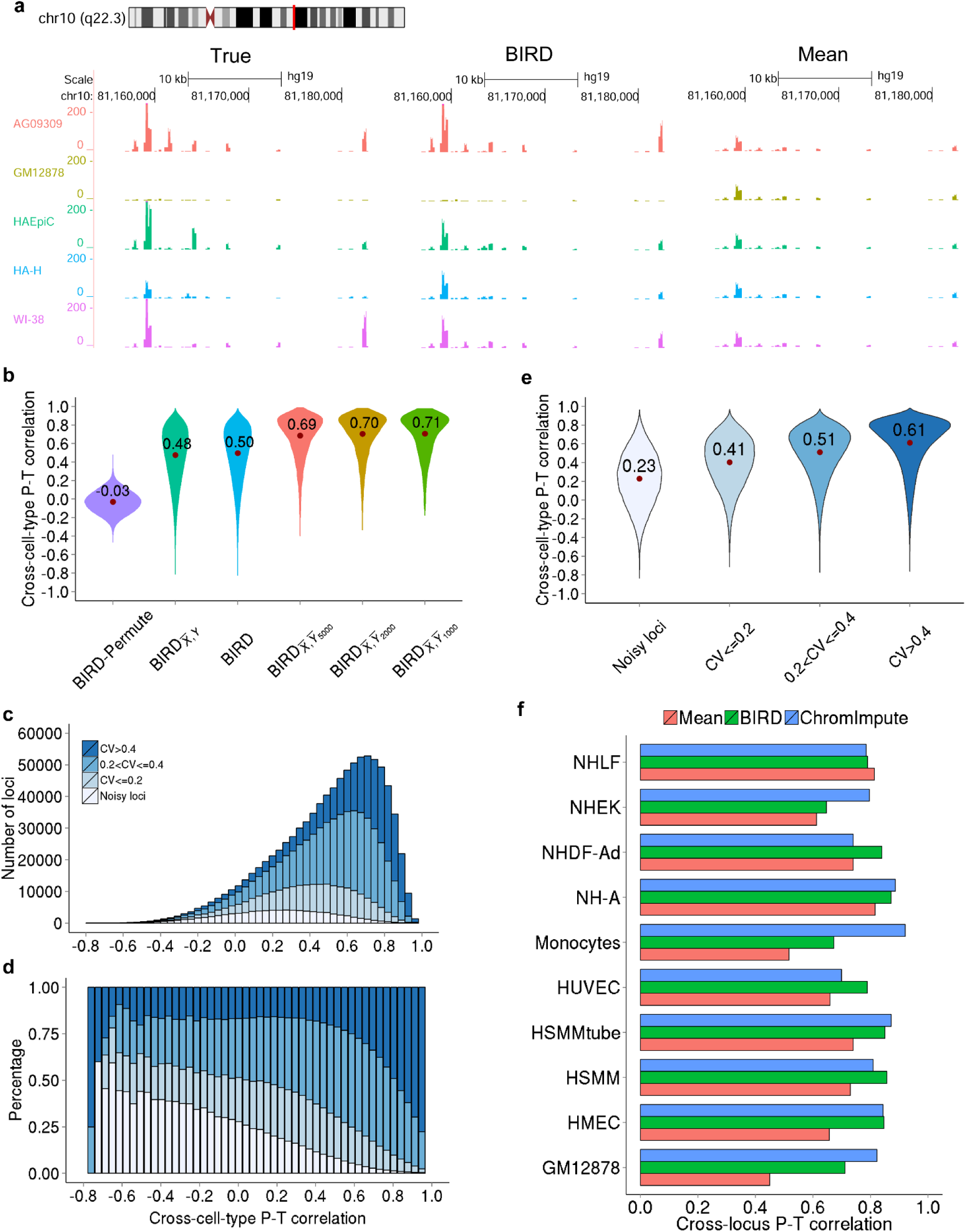
Cross-cell-type prediction performance and a comparison with ChromImpute. (**a**) An example of true and predicted DNase-seq signals for five different cell types. “True”: true DNase-seq bin read count; “BIRD”: DH signal predicted by BIRD; “Mean”: the average DH signal of training cell types. For “BIRD” and “Mean”, signals are transformed back from the log-scale to the original scale. (**b**) Comparison between locus-level and cluster-level predictions in terms of cross-cell-type prediction accuracy. For each method, distribution and mean of *r_C_* of all genomic loci or pathways (i.e., DHS clusters) are shown. DHSs were clustered into 1000, 2000 and 5000 clusters in 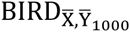, 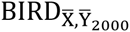 and 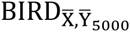 respectively. (**c**) Locus-level cross-cell-type prediction accuracy by BIRD. Genomic loci were grouped into four categories based on the coefficient of variation (CV) of the predicted DH values in 17 test cell types. The histogram shows the distribution of *r_C_* of all loci, stratified based on the four CV categories. (**d**) Loci are grouped into bins based on the cross-cell-type prediction accuracy *r_C_*. For each *r_C_* bin, the percentage of loci in each CV category is shown. (**e**) Distribution of *r_C_* for loci in each CV category. (**f**) Comparison between BIRD and ChromImpute. Cross-locus P-T correlation *r_L_* in 10 test cell types analyzed by both methods are shown. As a baseline, predictions based on the mean DH profile of training cell types are also shown.

*Cross-cell-type DH variation of regulatory element pathways can be predicted with substantially higher accuracy than that of individual loci*. This can be illustrated by comparing 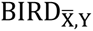 with 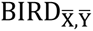. In 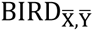, we first grouped correlated genomic loci into 1,000 clusters using the training data (**Methods**). Loci within each cluster share similar cross-cell-type DH variation pattern and hence can be viewed as a “pathway” consisting of co-activated regulatory elements (Thurman et al. 2012; Sheffield et al. 2013). 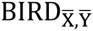 predicted the mean DH level of each cluster in each test cell type. The cross-cell-type P-T correlation *r_C_* for the cluster-level prediction was substantially higher than *r_C_* for the locus-level prediction (**Fig. 2b**, mean *r_C_* for 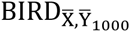 vs. 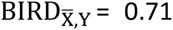 vs. 0.48). When genomic loci were grouped into 2,000 or 5,000 clusters, we obtained similar results (**Fig. 2b**). Thus, similar to gene set analysis (Subramanian et al. 2005), the overall cross-cell-type activity of a pathway of co-activated regulatory elements can be more reliably studied through prediction than that of individual loci. From a statistical perspective, the cluster mean can reduce the variance of random noise by averaging many measurements. Therefore, it can provide a cleaner signal that is easier to predict.

*The compound BIRD model improves locus-level prediction*. The compound BIRD model (denoted as BIRD) combines the locus-level prediction by 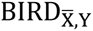 and cluster-level prediction by 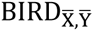 to balance the prediction bias and variance. As a result, it increased the locus-level prediction accuracy compared to 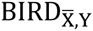 (**Fig. 1d-g, Methods**).

*Cross-cell-type prediction accuracy varies greatly among different loci*. For the compound BIRD model, *r_C_* of different genomic loci varied substantially (**Fig. 1e**, mean = 0.5). For 6% of loci, *r_C_* < 0 (i.e., prediction did not help). On the other hand, 56% and 20% of loci had *r_C_* > 0.5 and >0.75 respectively, indicating that DH could be predicted with moderate to high accuracy for a substantial fraction of loci. For each locus, we computed the coefficient of variation (CV) to characterize the variability of the predicted DH across the test cell types (**Methods**). We found that loci with poor cross-cell-type prediction accuracy (i.e., small *r_C_*) also tend to be less variable (i.e., had small CV) in the test cell types (**Fig. 2c,d**). Computing CV using the true DNase-seq data instead of the predicted DH yielded qualitatively similar results (**Supplementary Fig. 6**). One possible explanation for this phenomenon is that, compared to a highly variable locus, DH variation observed at a lowly variable locus is more likely due to random noise rather than true biological signals, and the correlation between predictions and random noise is expected to be zero. The CV of predicted DH provides a way to screen for loci whose cross-cell-type prediction is likely to be accurate. For instance, if we were to focus on loci with CV>0.4 rather than all loci, the mean *r_C_* would increase from 0.5 to 0.61, and 74% and 37% of loci would have *r_C_* > 0.5 and >0.75 respectively (**Fig. 2e**). Compared to the locus-level prediction by BIRD, cross-cell-type prediction accuracy by 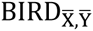 at the cluster-level was both more accurate and less variable (**Fig. 2b**). For 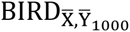, 84% and 55% clusters had *r_C_* > 0.5 and >0.75 respectively. The results were similar for 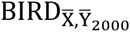 and 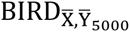.

*Comparisons of BIRD and ChromImpute*. ChromImpute is a recently developed method for imputing one functional genomic data type using multiple other data types (Ernst and Kellis 2015). We compared DH predictions by BIRD using only gene expression data with DH predictions by ChromImpute using multiple functional genomic data types (**Supplementary Methods**). Among 10 tested cell types, BIRD and ChromImpute showed comparable prediction performance. Neither method consistently outperformed the other (**Fig. 2f, Supplementary Fig. 7**). However, ChromImpute used ChIP-seq data for multiple histone modifications as predictors (these are the best predictors selected by ChromImpute for imputing DH (Ernst and Kellis 2015)), which are non-trivial to generate. By contrast, BIRD was based on gene expression data alone which are easier to generate and widely available.

*Robustness analysis*. The conclusions above do not depend on how the 57 cell types are partitioned into the training and testing data. We repeated the same analyses on four other random partitions (**Methods, Supplementary Table 1**), and similar results were obtained. For instance, **Supplementary Figure 8** shows that *r_L_*, *r_C_* and *τ* for BIRD from different partitions were similar.

### Predicting Transcription Factor Binding Sites Based on Gene Expression

We asked whether the predicted DH at DNA motif sites can predict transcription factor binding sites (TFBSs). Using BIRD models based on the 40 training cell types, we predicted TFBSs for 9 TFs in GM12878 cell line which was not in the training data. The predictions were evaluated using the corresponding TF ChIP-seq data from ENCODE in the same cell line. As a comparison, we also predicted TFBSs using true DNase-seq data (positive control) and using the mean DH profile of the training cell types (negative control). **Figure 3a-b** and **Supplementary Figure 9a-g** show how sensitivity of detecting motif-containing ChIP-seq binding sites changed with increasing number of predictions. For example, for TF ELF1 in GM12878, top 15000 BIRD (UW) predictions gave a sensitivity of 0.76 at an estimated false discovery rate (FDR, measured using *q*-value) of 0.05 (**Fig. 3b, Supplementary Methods**). As expected, true DNase-seq data predicted TFBSs better than BIRD. However, BIRD substantially improved the prediction based on the mean DH profile. For BIRD, predictions were made using exon array data generated by three different laboratories. The lab difference turned out to be smaller than the differences between prediction methods (**Fig. 3a-b, Supplementary Fig. 9a-g**). A similar analysis for 3 other TFs in K562 cell line yielded similar results (**Supplementary Methods, Fig. 3c**, **Supplementary Fig. 9h-i**).

**Figure 3.**
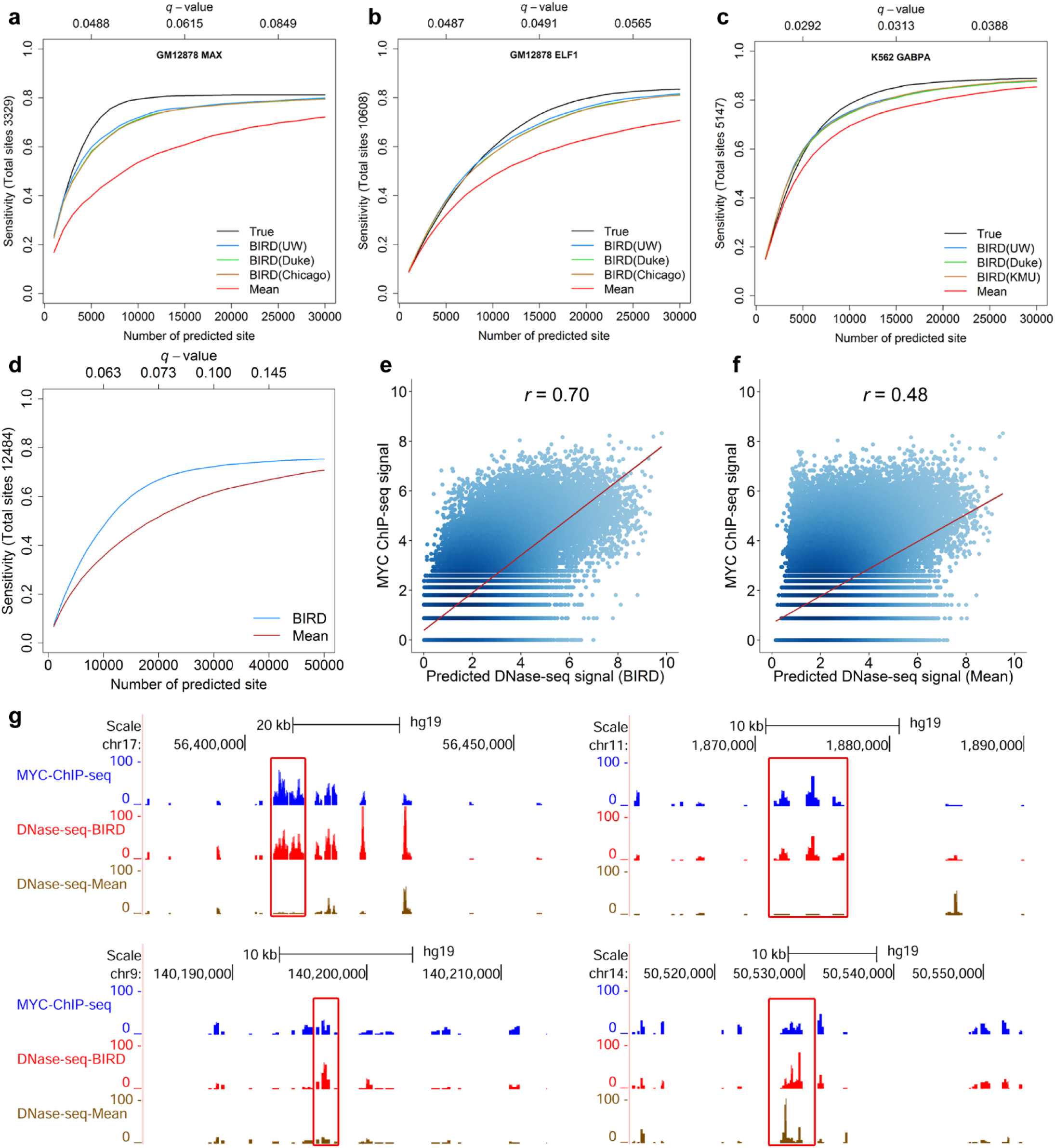
Predicting transcription factor binding sites. (**a**)-(**b**) Sensitivity-rank curve for predicting MAX and ELF1 binding sites in GM12878 using three different methods: true DNase-seq data (“True”), BIRD, and mean DH profile of training cell types (“Mean”). For BIRD, “BIRD(UW)”, “BIRD(Duke)” and “BIRD(Chicago)” denote predictions made using exon arrays generated by three different labs. For each method, the sensitivity-rank curve shows the percentage of true TFBSs that were discovered by top predicted motif sites. The *q*-values corresponding to top 5000, 15000, and 25000 predictions are shown on top of each plot. *q*-values from BIRD(UW) are shown, and *q*-values from the other two labs were similar and therefore not displayed for clarity. (**c**) Sensitivity-rank curve for predicting GABPA binding sites in K562. BIRD predictions were generated using exon arrays from three different labs. *q*-values from BIRD(UW) are shown. (**d**) Sensitivity-rank curve for predicting MYC binding sites in P493-6 using BIRD and the mean DH profile of training cell types. *q*-values from BIRD are shown. (**e**) True MYC ChIP-seq signal (log2 read count) in P493-6 versus BIRD predicted DH at all E-box motif sites. The correlation was high (*r_L_*=0.70). (**f**) True MYC ChIP-seq signal in P493-6 versus mean DH of training cell types at all E-box motif sites. The correlation was low (*r_L_*=0.48). (**g**) Examples showing true MYC ChIP-seq signal (read count, blue) in P493-6 and the predicted DH signal by BIRD (red) and “Mean” (brown). BIRD more accurately captured the true signal than “Mean” (highlighted with red boxes).

To further demonstrate BIRD in a realistic setting, we retrained BIRD using all 57 cell types for 1,108,603 loci with DH signal in at least one cell type. We then applied it to exon array data for P493-6 B cell lymphoma (a non-ENCODE cell line) generated by a non-ENCODE lab (Ji et al. 2011). We predicted MYC binding sites by identifying and ranking E-box motif sites CACGTG based on the predicted DH signal (**Supplementary Methods**). The predictions were evaluated using MYC ChIP-seq data (Sabò et al. 2014) in P493-6 cells (**Supplementary Methods**), from which 12,484 MYC binding peaks (FDR<0.01) overlapping E-box motif sites were discovered and served as the gold standard. **Figure 3d** shows the prediction performance. Among the top 20,000 predicted MYC binding sites (*q*-value < 0.073), 10,866 (54%) were indeed bound by MYC according to MYC ChIP-seq. The remaining 46% may represent a mixture of noise and true binding sites of other TFs since the E-box motif can also be recognized by multiple other TFs. In terms of sensitivity, 8,338 (67%) MYC binding peaks were overlapped with the predicted MYC binding sites (one peak may overlap with >1 DHSs). Thus, despite the fact that the training and test data have different lab origins, one can discover a substantial fraction of true MYC binding sites. The predicted DH also showed strong correlation with the true ChIP-seq signal (**Fig. 3e,g**). By contrast, predictions based on the mean DH profile of the 57 training cell types had substantially lower prediction accuracy (**Fig. 3d,f-g**). This demonstrates that in the absence of ChIP-seq data, one may use gene expression to predict TFBSs to identify promising follow-up targets.

### Regulome Prediction Based on 2000 Public Gene Expression Samples in GEO

The vast amounts of gene expression data from diverse biological contexts in GEO represent a resource that no single laboratory can generate. As a proof-of-principle test, we collected 2,000 human exon array samples from GEO and applied BIRD trained using all 57 ENCODE cell types for 1,108,603 loci to these samples to predict regulome. These predictions are made available as a web resource PDDB (<u>P</u>redicted <u>D</u>Nase I hypersensitivity <u>d</u>ata<u>b</u>ase). A user interface is provided for data query, display and download (**Fig. 4a-c, Methods, Supplementary Methods**).

**Figure 4.**
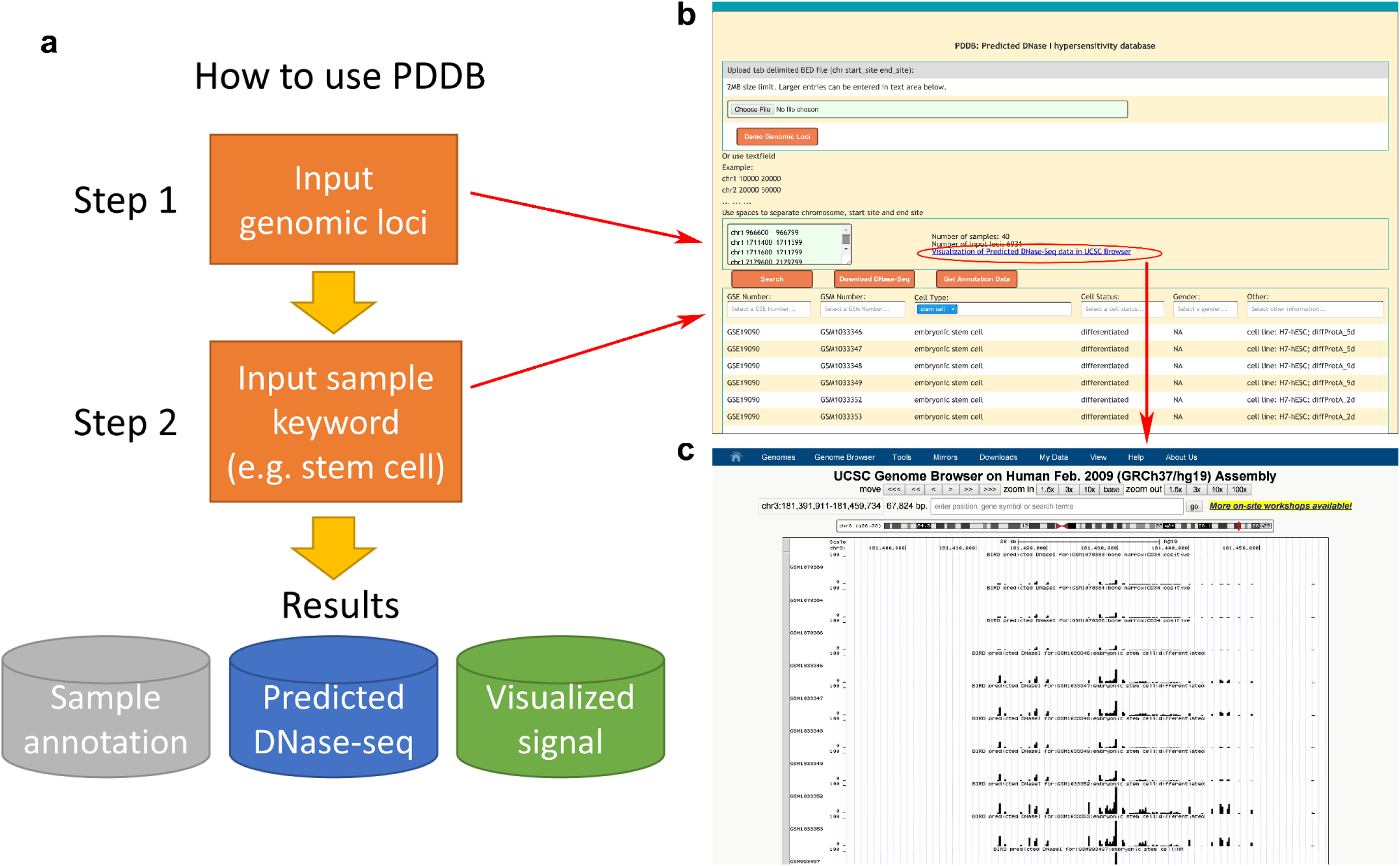
The predicted DNase I hypersensitivity database (PDDB). (**a**) Flowchart illustrating how to use PDDB. Step 1: provide a list of genomic loci of interest. Step 2: provide keywords in one or multiple annotation fields (e.g., type “stem cell” in the “Cell Type” column) to search for samples of interest. PDDB will return predicted DH for the queried loci and samples along with sample annotation and data for visualization. (**b**) Web interface of PDDB. Users can download the predicted DH data by clicking the “Download DNase-seq” button. The sample annotation data can be downloaded by clicking the “Get Annotation Data” button. (**c**) By clicking the “Visualization of Predicted DNase-Seq data in UCSC Browser” link in the PDDB web interface (red circle in **b**), one can display the predicted DH signal in the UCSC genome browser.

Researchers can use PDDB to explore regulatory element activities in biological contexts for which they do not have available regulome data. As a feasibility test, we first queried predicted DH for three genes FBL, LIN28A and BLMH in P493-6 B cell lymphoma (for which no public DNase-seq data are available) and H9 human embryonic stem cells. Promoters of these genes are known to be bound by MYC in a cell type dependent fashion (Ji et al. 2011). FBL is bound in both P493-6 and H9, LIN28A is bound in H9 but not in P493-6, and BLMH is bound in P493-6 but not in H9 (Koh et al. 2011; Chang et al. 2009; Ji et al. 2011). PDDB successfully predicted these known cell-type-dependent binding patterns (**Fig. 5a-c, Supplementary Fig. 10**).

**Figure 5.**
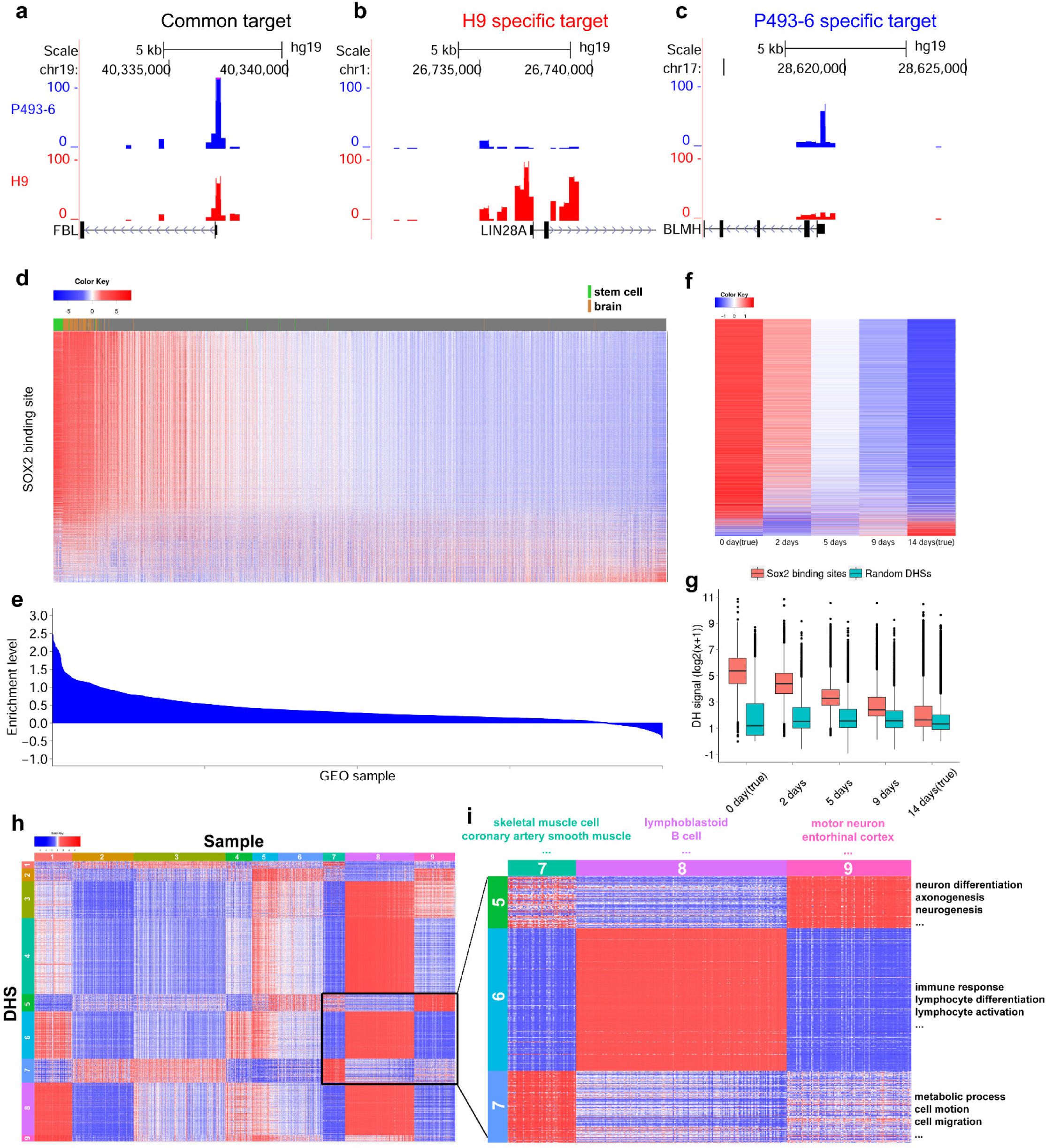
Predicting regulome using PDDB. (**a**)-(**c**) Predicted DH in promoter regions of FBL (**a**), LIN28A (**b**) and BLMH (**c**) in P493-6 B cell lymphoma and H9 embryonic stem cells. For H9, PDDB contains multiple replicate samples which produced similar results. One replicate is shown here and the other replicates are shown in **Supplementary Fig. 10**. (**d**) Predicted DH at SOX2 binding sites in 2,000 PDDB samples. Each column is a sample, and each row is a binding site. Values within each row are standardized to have zero mean and unit SD before visualization. (**e**) Relative DH enrichment level when comparing SOX2 binding sites with random sites (**Supplementary Methods**). (**f**) Predicted DH at SOX2 binding sites in H7 stem cells after 2, 5 and 9 days of differentiation. True DH from undifferentiated H7 cells and cells at differentiating day 14 in the training data are also shown. Rows are SOX2 sites and columns are time points. Values within each row are standardized before visualization. (**g**) Predicted DH at SOX2 binding sites are compared with predicted DH at random DHSs. At each time point, DH values from all sites are displayed using a boxplot. (**h**) Predicted DH at 2,011 MEF2A binding sites in 1,061 MEF2A-expressing PDDB samples (**Supplementary Methods**). Each column is a sample. Each row is a MEF2A binding site. Values within each row are standardized before visualization. Samples and DHSs were clustered. (**i**) The highlighted region in (**h**) which shows DHS-clusters with increased DH in muscle, lymphoblastoid, and brain related samples, respectively.

Next, we obtained a list of SOX2 binding sites in human embryonic stem cells from a published ChIP-seq study (Watanabe et al. 2014) (**Supplementary Methods**). **Figure 5d** shows the predicted DH at these sites across the 2,000 GEO samples. The samples were ordered based on the overall DH enrichment level at all SOX2 binding sites relative to random genomic sites (**Supplementary Methods**, **Fig. 5e**). Samples with strong predicted DH at SOX2 binding sites include stem cells (green bar in **Fig. 5d**) and brain (brown bar), consistent with known roles of SOX2 in these sample types (Chambers and Tomlinson 2009; Takahashi and Yamanaka 2006; Ferri et al. 2004; Phi et al. 2008). Interestingly, PDDB contained differentiating H7 embryonic stem cells collected at day 2, 5 and 9 after initiation of differentiation. Our 57 training cell types contained undifferentiated H7 cells and H7 cells at differentiating day 14. Together, these samples formed a time course. Examination of the predicted DH for day 2, 5, and 9 along with the true DH for day 0 and 14 shows that the predicted DH at SOX2 binding sites decreased as the differentiation progressed (**Fig. 5f-g**), consistent with the known role of SOX2 for maintaining the undifferentiated status of stem cells (Takahashi and Yamanaka 2006; Chambers and Tomlinson 2009). Thus, the dynamic changes of SOX2 binding activities were correctly predicted in PDDB.

The above examples show that expression samples in GEO can be used to meaningfully predict DH. With ChIP-seq data for a TF from one biological context, one may also use PDDB to systematically explore in what other biological contexts each binding site might be active, and group TFBSs into functionally related subclasses accordingly. For instance, we obtained MEF2A ChIP-seq binding sites in GM12878 lymphoblastoid cells from ENCODE. MEF2A is a TF involved in muscle development (Edmondson et al. 1994) and neuronal differentiation (Flavell et al. 2008). Using PDDB (**Supplementary Methods**, **Fig. 5h-i**, **Supplementary Fig. 11, Supplementary Tables 5-6**), we first clustered samples and MEF2A binding sites into different groups and performed functional annotation analysis on each group using the Database for Annotation, Visualization and Integrated Discovery (DAVID) (Huang et al. 2009; Huang et al. 2008). A group of MEF2A binding sites associated with genes involved in cell motion, cell migration and regulation of metabolic processes was found to be more active in muscle related samples (including coronary artery smooth muscle and cardiac precursor cell which are not covered by ENCODE) than in lymphoblastoid (**Fig. 5h-i**). Another group of sites associated with neuron differentiation and neurogenesis genes was found to be more active in neuron and brain related samples (including non-ENCODE sample types such as entorhinal cortex and motor neuron) (**Fig. 5h-i**). This demonstrates how PDDB can provide a more detailed view of TFBSs not offered by the original experiment in GM12878, and how PDDB can be used to investigate many biological contexts not covered by ENCODE.

### Predictions as Pseudo-Replicates to Improve Analyses of DNase-seq and ChIP-seq Data

In applications of high-throughput regulome profiling technologies, it is common to encounter data with low signal-to-noise ratio or small replicate number. Both can lead to low signal detection power. However, if one has gene expression data, BIRD predictions may be used as pseudo-replicates to enhance the signal. As a test, we analyzed DNase-seq data for GM12878 generated by ENCODE. The data had two replicates. We reserved one replicate as “truth” and used the other one as the “observed” data. Applying the BIRD prediction models trained earlier using the 40 training cell types (GM12878 not included), we predicted DH in GM12878 and treated the prediction as a pseudo-replicate. We then estimated “true” DH using either the “observed” data alone (obs-only) or the average of the “observed” data and pseudo-replicate (BIRD+obs). After adding the pseudo-replicate, the correlation between the predicted and true DH increased (**Fig. 6a-b**, *r_L_* for BIRD+obs vs. obs-only = 0.82 vs. 0.77). Replacing BIRD predictions with the mean DH profile of 40 training cell types in this analysis (Mean+obs) did not yield similar increase in the P-T correlation (*r_L_*= 0.76). We carried out the same analyses on 16 test cell types, and BIRD predictions improved signal in 12 of them (**Fig. 6c, Supplementary Methods**).

**Figure 6.**
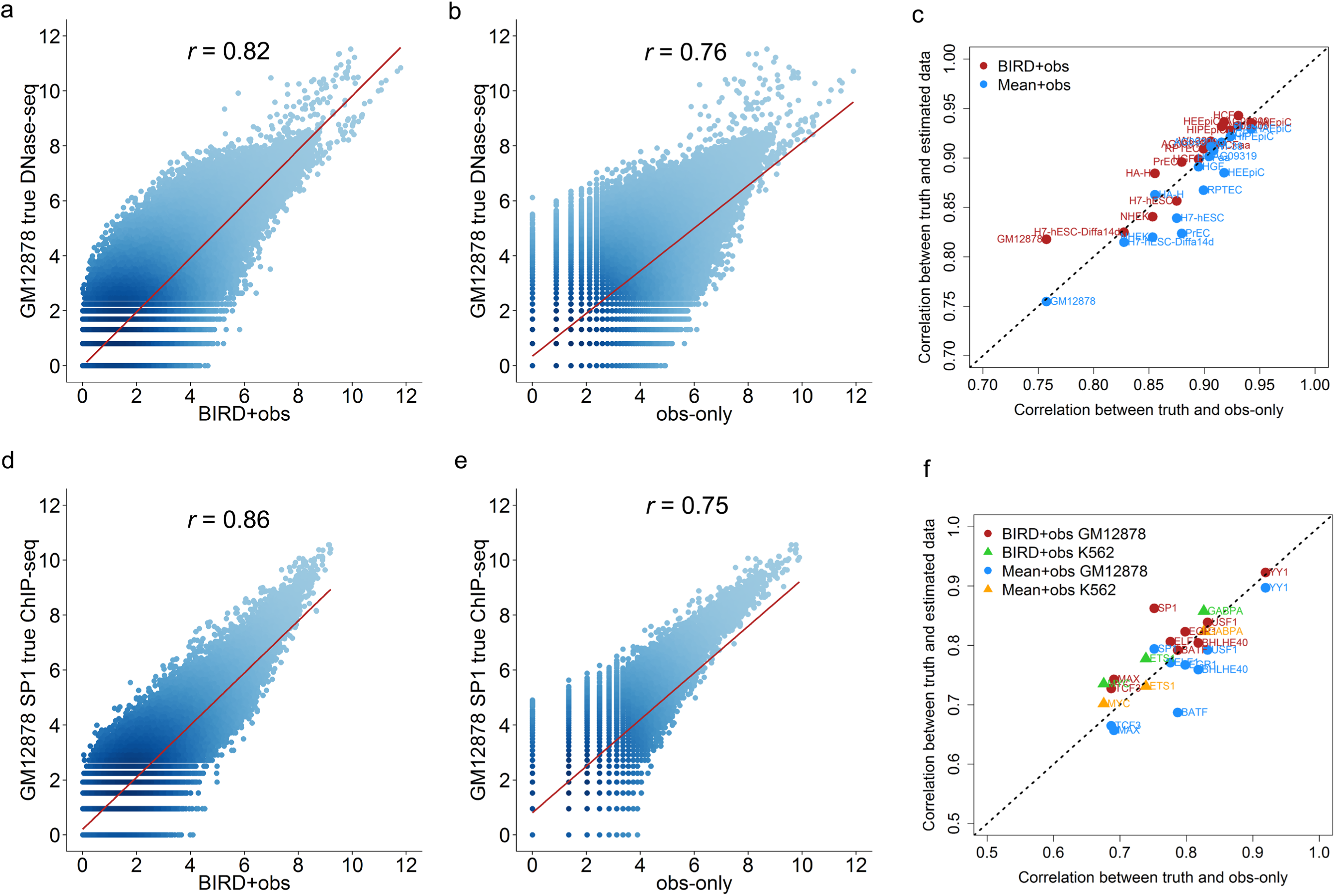
BIRD predictions used as pseudo-replicates to improve DNase-seq and ChIP-seq data analyses. The observed signal from one sample (“obs-only”) in a test cell type was combined with BIRD predictions to produce the integrated signal (“BIRD+obs”). Signals before and after integration are compared with the observed signal from another sample from the same cell type (“truth”). (**a**)-(**c**) DNase-seq. (**a**) Correlation (*r*) between the “truth” and “BIRD+obs” (i.e., the integrated signal) in GM12878. Each dot is a genomic locus. (**b**) Correlation between the “truth” and “obs-only” (i.e., the original signal without integrating BIRD) in GM12878. (**c**) The same analyses were done for 16 test cell types. Red dots in the scatterplot compares the P-T correlation *r* for BIRD+obs vs. *r* for obs-only in the 16 cell types. BIRD+obs outperformed obs-only in 12 of 16 test cell types. As a control, BIRD was replaced by the mean DH profile of training cell types. Blue dots show the P-T correlation *r* for Mean+obs vs. *r* for obs-only in the 16 test cell types. Mean+obs did not improve over obs-only. (**d**)-(**f**) ChIP-seq. (**d**) Correlation between the “truth” and “BIRD+obs” for SP1 in GM12878 at SP1 motif sites. (**e**) Correlation between the “truth” and “obs-only” for SP1 in GM12878 at SP1 motif sites. (**f**) The same analyses were done for 9 TFs in GM12878 (circles) and 3 TFs in K562 (triangles). Once again, BIRD+obs outperformed obs-only in 11 of 12 cases (red and green), but Mean+obs did not improve over obs-only (blue and yellow)

Similarly, we tested if the predicted DH can boost ChIP-seq signals using ChIP-seq data for 9 TFs in GM12878 and 3 TFs in K562 (**Supplementary Methods**). Similar results were observed (**Fig. 6d-f**). BIRD+obs outperformed obs-only in nearly all test cases (11 out of 12 TFs). Together, these results show that predictions can serve as a bridge to integrate expression and regulome data so that one can more effectively use available information to improve data analysis.

## DISCUSSION

In summary, this study for the first time examined systematically to what extent regulatory element activities can be predicted by gene expression alone. We developed BIRD for big data prediction. The study also demonstrates the feasibility of using gene expression to predict TFBSs, applying BIRD to GEO to expand the current regulome catalog, and using predictions to facilitate data integration. BIRD is a novel approach to extract information from gene expression data to study regulome. In the absence of experimental regulome data (e.g., ChIP-seq or DNase-seq data), BIRD predictions can provide valuable information to guide hypothesis generation, target prioritization, and design of follow-up experiments. When experimental regulome data are available, BIRD predictions can also serve as pseudo-replicate to improve the data analysis. In a companion study, we show that BIRD can also predict DH using RNA-seq and in samples with small number of cells, and it can outperform state-of-the-art technologies for mapping regulome in small-cell-number samples (Zhou et al. submitted).

Our results have important practical implications for the analysis of existing and future gene expression data. Conventionally, gene expression data are mainly collected to study transcriptome. The method and software developed in this study now allow one to conveniently utilize such data to study gene regulation. By adding a new component to the standard analysis pipeline of expression data, expression-based regulome prediction can bring added value to an enormous number of new and existing gene expression experiments. Given the wide application of gene expression profiling, this will greatly impact how expression data are most effectively used.

Compared to conventional regulome mapping technologies, BIRD also has its unique advantages. Since gene expression profiling experiments are more widely conducted than regulome mapping experiments, the number of biological contexts with gene expression data is orders of magnitude larger than the number of contexts with experimental regulome data. BIRD can be readily applied to massive amounts of existing and new gene expression data to generate regulome information for a large number of biological contexts without experimental regulome data. In the near future, no other experimental regulome mapping technology can achieve similar level of comprehensiveness in terms of biological context coverage.

Our current study may be extended in multiple directions in the future. For instance, it is important to extend BIRD to other gene expression platforms. It also remains to be answered whether gene expression can be similarly used to predict other functional genomic data types.

## METHODS

### DNase-seq data processing

The bowtie (Langmead et al. 2009) aligned (alignment based on hg19) DNase-seq data for 57 human cell types with normal karyotype were downloaded from the ENCODE in bam format (download link: http://hgdownload.cse.ucsc.edu/goldenPath/hg19/encodeDCC/wgEncodeUwDnase). The human genome was divided into base pair (bp) non-overlapping bins. The number of reads falling into each bin was counted for each DNase-seq sample. To adjust for different sequencing depths, bin read counts for each sample *i* were first divided by the sample’s total read count *N_i_* and then scaled by multiplying a constant 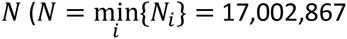, which is the minimum sample read count of all samples). After this procedure, the raw read count *n_li_* for bin *l* and sample *i* was converted into a normalized read count 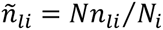. The normalized read counts from replicate samples were averaged to characterize the DH level for each bin in each cell type. The DH level was then log2 transformed after adding a pseudocount 1. The transformed data were used for training and testing prediction models, treating each bin as a genomic locus. Since chromosome Y was not present in all samples, we excluded this chromosome from our subsequent analyses.

### Gene expression data processing

The Affymetrix Human Exon 1.0 ST Array (i.e. exon array) data for the same 57 ENCODE cell types were downloaded from GEO (GEO accession number: GSE19090). Additionally, we downloaded 2000 exon array samples from GEO for constructing the PDDB database (GEO accession numbers for these samples are available at PDDB). All samples were processed using the GeneBASE (Kapur et al. 2007) software to compute gene-level expression. The output of GeneBASE was expression levels of 18,524 genes in each sample. The GeneBASE gene expression levels were log2 transformed after adding a pseudocount 1 and then quantile normalized (Bolstad 2015) across samples. For the 57 ENCODE cell types, replicate samples within each cell type were averaged and the averaged mean expression profile of each cell type was used for training and testing the prediction models.

### Training-test data partitioning and genomic loci filtering

The 57 ENCODE cell types were randomly partitioned into a training dataset with 40 cell types and a test dataset with 17 cell types (**Supplementary Table 1**, partition # 1). Since not all genomic loci are regulatory elements, we first screened for genomic loci with unambiguous DH signal in at least one cell type in the training data as follows. Genomic bins with normalized read count >10 in at least one cell type were identified and retained, and the other genomic bins were excluded. Among the retained loci, bins with normalized read count >10,000 in any cell type were considered abnormal and these bins were also excluded from subsequent analyses. Finally, for each remaining bin, a signal-to-noise ratio (SNR) was computed in each cell type, and bins with small SNR in all cell types were filtered out. To compute SNR of a genomic bin in a cell type, we first collected 500 bins in the neighborhood of the bin in question. Then, we computed the average DH level of these bins. Next, the DH level was log2 transformed after adding a pseudocount 1 to serve as the background. The log_2_(SNR) was defined as the difference between the normalized and log2 transformed DH level of the bin in question and the background. Genomic bins with log_2_(SNR)>2 in at least one cell type were identified and retained for subsequent analyses, and the other genomic bins were excluded. After applying this filtering procedure to the 40 training cell types, 912,886 genomic bins were retained and used for training and testing prediction models in **Figures 1** and **2**. Bins selected by this procedure were referred to as DNase I hypersensitive sites (DHSs) in this article. We note that the above filtering procedure only uses the training cell types. This allows one to objectively evaluate the prediction performance in real applications where models trained using the training cell types are applied to make predictions in new cell types for which DNase-seq data are not available.

In order to evaluate the robustness of our conclusions, we repeated the same random partitioning procedure five times, resulting in five different training-test data partitions (**Supplementary Table 1**). For each partition, genomic loci were filtered using the same protocol described above, and the retained loci (which depend on the training data and therefore are different for different partitions) were used to train and test BIRD. Results from the first partition were presented in the main article, and results from the other four random partitions were similar (**Supplementary Fig. 8**).

For predicting TFBSs in K562 and P493-6 B cell lymphoma and the analyses of 2000 GEO exon array samples used for constructing PDDB, prediction models were retrained using all 57 ENCODE cell types as training data. Applying the genomic loci filtering protocol described above to these 57 cell types resulted in 1,108,603 genomic bins for which prediction models were constructed and evaluated.

### Notations and problem formulation

For a biological sample, let *Y_l_* be the DH level of genomic locus *l*(=1,…,*L*), and let *X_g_* be the expression level of gene *g* (=1,…,*G*). The genome-wide DH profile and gene expression profile are represented by two vectors ***Y*** = (*Y*_1_,…,*Y_L_*)*^T^* and ***X*** = (*X*_1_,…,*X_G_*)*^T^* respectively. Here, the superscript *T* indicates matrix or vector transpose. Both the DH and gene expression profiles are assumed to be normalized and at log2 scale. Our goal is to use ***X*** to predict ***Y***. This can be formulated as a problem of building a regression ***Y****_l_* = *f_l_*(*X*) + *ϵ_l_* for each genomic locus. Here *ϵ_l_* represents random noise, and *f_l_*(.) is the function that describes the systematic relationship between the DH level of locus *l* (i.e., *Y_l_*) and the gene expression profile (i.e., ***X***).

The function *f_l_*(***X***) is unknown. We train it using ***X*** and ***Y*** observed from a number of different cell types. The training data are organized into two matrices: a gene expression matrix 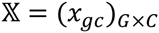 and a DH matrix 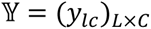. Rows in these matrices are genes and genomic loci respectively. Columns in these matrices are cell types. *C* is the number of training cell types. Each column of 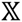 and 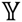 is a realization of the random vector ***X*** and ***Y*** in a specific cell type. Building the prediction model for each locus *l* is a challenging high-dimensional regression problem since the dimensionality of the predictor ***X*** is much bigger than the sample size of the training data (i.e., *G* ≫ *C*). What makes this problem even more challenging than the conventional high-dimensional problems in statistics is that one needs to solve a massive number of such high-dimensional regression problems (one for each locus) simultaneously. Thus it is important to consider both statistical efficiency and computational efficiency when developing solutions.

In subsequent sections, various methods for training *f_l_*(***X***) will be described. Each method has a training component and a prediction component. Before training prediction models, we standardize each row of 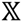 and 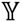 in the training data to have zero mean and unit standard deviation (SD). More precisely, each DH value in 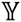 is standardized using 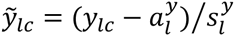 where 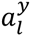 and 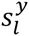 are the mean and SD of the DH signals at locus *l* (i.e., row *l* of 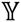). Similarly, each expression value in 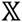 is standardized using 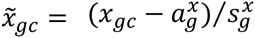 where 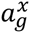 and 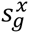 are the mean and SD of the gene expression for gene *g* (i.e., row *g* of 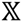). The prediction models are then constructed using the standardized values 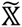 and 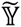.

Once the models are constructed using the training data, they can be applied to new samples to make predictions. To do so, the expression profile ***X*** of the new sample is first quantile normalized to the quantiles of the training exon array data. The log2-transformed expression value of each gene *X_g_* in the new sample is then standardized using 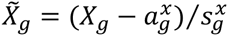, where 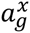 and 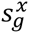 are the pre-computed mean and SD of the gene expression for gene *g* in the training data. After applying the trained model to the standardized gene expression profile 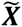 to make predictions, the predicted DH value for each locus, 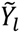, is transformed back using 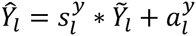, where 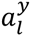 and 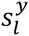 are the pre-computed mean and SD of the DH signals for locus *l* in the training data. The unstandardized 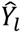 gives the prediction for *Y_l_*, the DH level of genomic locus *l* in the new sample.

### Measures for method evaluation

In order to evaluate prediction performance of a prediction method, the method can be applied to a number of test cell types to predict their DH profiles based on their gene expression profiles. Let 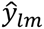 be the predicted DH level of locus *l* in test cell type *m* (=1,…,*M*), and let *y_lm_* be the true DH level measured by DNase-seq (both are at log2 scale). Three performance statistics were used in this study (**Fig. 1c**):

1. Cross-locus correlation (*r_L_*). This is the Pearson’s correlation between the predicted signals 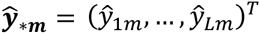 and the true signals 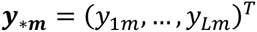 across different loci for each test cell type *m*. The cross-locus correlation measures the extent to which the DH signal within each cell type can be predicted.
2. Cross-cell-type correlation (*r_C_*). This is the Pearson’s correlation between the predicted signals 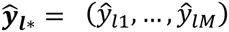 and the true signals ***y****_l*_*=(*y_l_*_1_,…,*y_lM_*) across different cell types for each locus *l*. The cross-cell-type correlation measures how much of the DH variation across cell type can be predicted.
3. Squared prediction error (*τ*). This is measured by the total squared prediction error scaled by the total DH data variance in the test dataset: 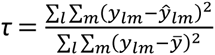, where 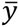 is the mean of *y_lm_* across all DHSs and test cell types.

### Prediction based on neighboring genes

For each genomic locus *l*, *N* closest genes were identified (gene annotation based on RefSeq genes of human genome hg19 downloaded from UCSC genome browser: http://hgdownload.cse.ucsc.edu/501goldenPath/hg19/database/refFlat.txt.gz). The closeness was defined by the distance between the gene’s transcription start site and the locus center. Using the selected genes 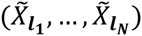 as predictors, a multiple linear regression 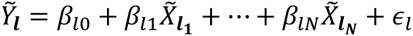 is fitted. Based on the fitted model, the standardized DH level of locus *l* in a new sample is predicted using 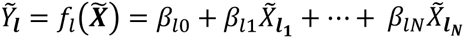. We tested different values of *N* (= 1, 2,…, 20) on a randomly selected set of DHSs (n=9,128; ~1% of the 912,886 DHSs obtained from the 40 training cell types). The performance for the neighboring gene approach shown in **Figure 1d-g** was based on the performance achieved at the optimal *N*. For instance, **Supplementary Figure 2a** shows the *r_C_* distribution for different *N* based on the 9,128 DHSs. At *N*=15, the mean *r_C_* reached its maximum. Correspondingly, the *r_C_* distribution shown in **Figure 1e** was based on *N*=15.

We also tested whether nonlinear regression can improve the prediction. Generalized additive model with smoothing spline (GAM) was applied (using R package “gam” (Hastie 2015)) to the same 1% of DHSs. However, the best prediction performance of GAM was worse than the best prediction performance of the linear regression (**Supplementary Fig. 2a**, see the best performance of GAM achieved at *N* = 17 vs. the best performance of linear model achieved at *N* = 15). This indicates that using non-linear model did not improve prediction accuracy. Moreover, the computational time required by GAM was substantially longer than linear regression (**Supplementary Fig. 2b**), making it difficult to apply to the whole genome. Based on this, linear regression was used to perform our genome-wide analysis.

### 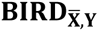 – The elementary BIRD model

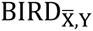 is the basic building block of BIRD. This approach first groups correlated genes into clusters. This is achieved by clustering rows of the standardized training data matrix 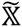 into *K* clusters using k-means clustering (Hartigan and Wong 1979) (Euclidean distance used as similarity measure). Based on the clustering result, the gene expression profile 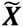 of each sample is converted into a lower dimensional vector 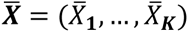, where 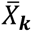 is the mean expression level of genes in cluster *k*. BIRD will use gene clusters’ mean expression 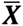 instead of the expression of individual genes 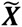 as predictors to build prediction models. Clustering serves multiple purposes. It reduces the dimension of the predictor space. By combining correlated genes, it also reduces the co-linearity among predictors. Additionally, cluster mean is less sensitive to measurement noise and therefore can reduce the impact of measurement error of a gene on the prediction.

After clustering, the *G* × *C* matrix 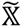 is converted into a *K* × *C* matrix 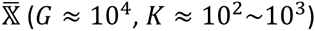. The predictor dimension is reduced, but it is still high compared to sample size. Borrowing the idea from the recent high-dimensional regression literature (Fan and Lv 2008), we further reduce the predictor dimension using a fast variable screening procedure: for each DHS locus *l*, the Pearson’s correlation between its DH signal (i.e., row *l* of 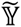) and the expression of each gene cluster *k* (i.e., row *k* of 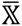) across the training cell types is computed, and the top *N* (≈ 10^1^) clusters with the largest correlation coefficients are selected. Using the selected clusters 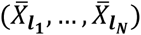 as predictors, a multiple linear regression 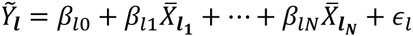 is then fitted. Based on the fitted model, the standardized DH level of locus *l* in a new sample is predicted by 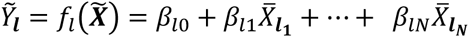. Of note, although each regression model only contains a small number of predictors, these predictors are selected after examining information from all genes. Therefore, training the prediction model utilizes information from all genes.

The elementary BIRD model has two parameters: the cluster number *K* and the predictor number *N*. In this study, we set *K*=1500 and *N*=7. These parameters were chosen based on testing different values of *K* and *N* (*K*=100, 200, 500, 1000, 1500, 2000; *N*=1, 2, 3, 4, 5, 6, 7, 8) using a 5-fold cross-validation conducted within the 40 training cell types (i.e., the same training cell types used for **Figs. 1** and **2**) on a random subset of genomic loci (1% of all DHSs). Since cross-cell-type prediction is more difficult than cross-locus prediction, we identified the optimal parameter combination as the one that maximizes the mean cross-cell-type correlation *r_C_*. **Supplementary Figure 3a** shows that the optimal combination was *K*=1500 and *N*=7. This parameter combination was then used in all subsequent 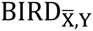, 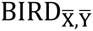, and compound BIRD models throughout this study.

In **Supplementary Methods and Supplementary Figure 4**, we compared the elementary BIRD model 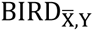 with a number of alternative prediction methods including Lasso (Tibshirani 1996), linear regression with stepwise predictor selection (Hocking 1976) (SPS), k-nearest neighbors (Altman 1992) (KNN) and random forest (Breiman 2001) (RF) using 1% of the DHSs obtained from the 40 training cell types. This benchmark analysis shows that the elementary BIRD model not only offers the best prediction accuracy but also is computationally efficient. Based on this result, 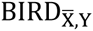 was used as the basic building block for subsequent modeling.

### BIRD_X,Y_ model

If one does not cluster co-expressed genes in the elementary BIRD model, 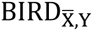 reduces to BIRD_X,Y_. In other words, BIRD_X,Y_ is a special case of 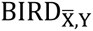 when the gene cluster number *K* is equal to the gene number *G*. BIRD_X,Y_ is not used in the final BIRD compound model. However, in **Figure 1d-f**, BIRD_X,Y_ and 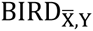 were compared to study the effect of gene clustering on prediction. BIRD_X,Y_ only has one parameter: the number of predictors *N*. Based on 5-fold cross-validation performed on the 40 training cell types using 1% of all DHSs from these training cell types, we identified *N* = 5 as the optimal value for BIRD_X,Y_ (**Supplementary Fig. 3a,b**). BIRD_X,Y_ based on this optimal *N* (*N* = 5) was compared to 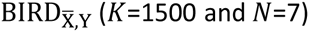 in **Figure 1d-g**. In **Supplementary Figure 3b**, BIRD_X,Y_ and 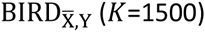 were also compared when both methods used the same *N*. In both comparisons, 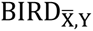 consistently outperformed BIRD_X,Y_.

### 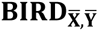 model

In addition to clustering co-expressed genes, 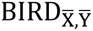 also groups genomic loci with similar DH patterns into clusters. This is done by clustering rows of the standardized matrix 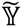 into *H* clusters using k-means clustering (Euclidean distance used as similarity measure). Based on the clustering result, the DH profile 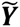 of each sample can be converted into a lower dimensional vector 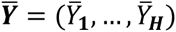, where 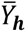 is the mean DH level of DHSs in cluster *h*. Instead of predicting the DH level 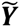 of individual loci, 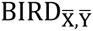 uses the cluster-level gene expression 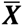 to predict cluster-level DH 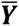. The prediction models are constructed using linear regression in a way similar to how the regression models are constructed in 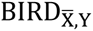. In **Figure 2b**, comparisons between 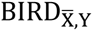 and 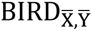 was used to illustrate cluster-level DH can be predicted with higher accuracy than DH at individual genomic loci. The same parameter combination *K*=1500 and *N*=7 was set for both 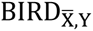 and 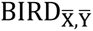. For 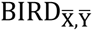, *H* was set to 1000, 2000 and 5000 respectively.

### BIRD – The compound BIRD model

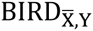 is a special case of 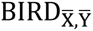 when DHSs are not clustered (i.e., *H* = *L*). Compared to 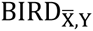, the increased accuracy of cluster-level prediction by 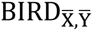 is partly because a cluster’s mean DH is usually associated with smaller variance of measurement noise than the DH level of individual loci. In 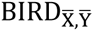, one may use the predicted cluster mean as the predicted DH level of each individual locus within the cluster. This will also generate a prediction for each locus. This locus-level prediction may be biased, but it is usually associated with smaller variance. By contrast, predictions by 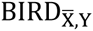 for each locus may be less biased but has larger variance. This motivates the compound BIRD model.

In the compound BIRD model, multiple 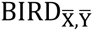 models with different *H* values are combined through model averaging, a useful technique to improve prediction accuracy by balancing the variance and bias. Consider making predictions for a sample. Let 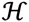 be the set of *H* values used by 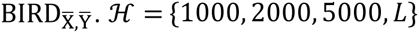 in this study. For each DHS locus *l*, let 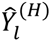 denote the locus-level DH predicted by 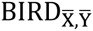 using cluster number 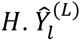 is the locus-level DH predicted by 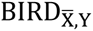. The compound BIRD model predicts the locus-level DH for locus *l* using a weighted average

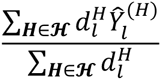

where 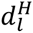 is the weight. For a given cluster number *H*, the weight 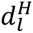 is determined using training data as follows. Let 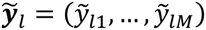 be the standardized locus-level DH for locus *l* observed in *M* training cell types. Each locus *l* is associated with a cluster. Let 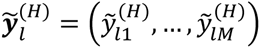 represent the average of the standardized DH level of all loci within the cluster corresponding to locus *l* in the *M* training cell types. Define 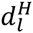 as the Pearson’s correlation between the two vectors 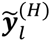 and 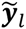. Note that when *H* = *L*, 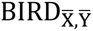 reduces to 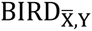, and we have 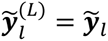 and 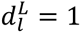. Thus the weight for 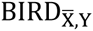 is 1.

Comparisons between the compound BIRD model (referred to as “BIRD”) and 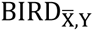 in **Figure 1d-g** show that BIRD outperforms 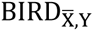. Therefore, the compound BIRD model was used as our final prediction model, and it was used for predicting TFBS, constructing PDDB, and improving DNase-seq and ChIP-seq data analyses.

### Random prediction models by permutation

To construct random prediction models, we permuted the cell type labels of DNase-seq data in the training dataset. This permutation broke the connection between DNase-seq and gene expression data. BIRD was then trained using the permuted training dataset, and the trained model was applied to predict DH in the test dataset. The permutation was performed 10 times. The prediction performance *r_L_*, *r_C_* and *τ* were computed for each permutation. The average values of these three statistics from the 10 permutations were used to represent the prediction performance of random prediction models.

### Wilcoxon signed-rank test for comparing different methods

In order to generate **Figure 1g**, two-sided Wilcoxon signed-rank test was performed to obtain *p*-values for comparing prediction accuracy of each pair of methods. For instance, in order to test whether two methods A and B perform equally in terms of *r_L_*, the paired *r_L_* values from these two methods for each cell type was obtained. Then the *r_L_* pairs from all cell types are used for the Wilcoxon signed-rank test. Similarly, to compare methods A and B in terms of *r_C_*, the paired *r_C_* values for each locus was obtained, and *r_C_* pairs from all genomic loci were used for the Wilcoxon signed-rank test.

### Categorization of test genomic loci when studying cross-cell-type correlation

When studying the cross-cell-type prediction performance (i.e., *r_C_*) of BIRD in **Figure 2c-e**, genomic loci were grouped into different categories based on their DH profile in the test cell types. First, because test cell types were not used to select genomic loci, a subset of selected genomic loci may not contain strong or meaningful DH signal in any test cell type. For such loci, the cross-cell-type correlation between the predicted and true DH signals (which are essentially noise) is expected to be low. For this reason, we identified DHSs with predicted DH level (log2 transformed) smaller than 2 in all 17 test cell types and labeled them as “noisy loci” (**Fig. 2c**). After excluding the noisy loci, the other loci were then categorized based on the coefficient of variation (CV) of the cross-cell-type DH values. For each locus, CV was calculated as the ratio of the standard deviation to mean of the predicted DH at this locus across all test cell types. Loci were divided into three categories: CV≤0.2, 0.2<CV≤0.4, CV>0.4 (**Fig. 2c**). A large CV indicates that the DH of a locus has more variation across cell types. **Figure 2c** shows the distribution of *r_C_*. Genomic loci are grouped into bins based on *r_C_* values. For each bin, the number of loci in different CV categories is shown. **Figure 2d** shows the percentage of loci in different CV categories for each *r_C_* bin. **Figure 2e** shows distribution of *r_C_* values for each CV category.

We also computed CV using the true DH values from the test DNase-seq data rather than predicted DH values. The results that loci with large *r_C_* also tend to have large CV remain qualitatively the same (**Supplementary Fig. 6**). In practice, however, since BIRD is typically used when DNase-seq data are not available, one can only use CV based on predicted DH values.

### The Predicted DNase I hypersensitivity database (PDDB)

PDDB is available at http://jilab.biostat.jhsph.edu/~bsherwo2/bird/index.php. Details on database construction and use are provided in **Supplementary Methods**.

### Software

BIRD software is available at https://github.com/WeiqiangZhou/BIRD. Models trained using the 57 ENCODE cell types have been stored in the software package. With these pre-compiled prediction models, making predictions on new samples provided by users is computationally fast. On a computer with 2.5 GHz CPU and 10Gb RAM, it took less than 2 minutes to make predictions for ~1 million DHSs in 100 samples.

### Other data analysis protocols

Procedures for comparing BIRD with other prediction methods, TFBS prediction, MYC, SOX2 and MEF2A analyses using PDDB, and improving DNase-seq and ChIP-seq signals are provided in **Supplementary Methods**.

## ACKNOWLEDGMENTS

The authors would like to thank Drs. X. Shirley Liu and Yingying Wei for insightful discussions. This research is supported by grants from the Maryland Stem Cell Research Fund (2012-MSCRFE-0135-00) and the National Institutes of Health (R01HG006282 and R01HG006841).

